# Mild and severe SARS-CoV-2 infection induces respiratory and intestinal microbiome changes in the K18-hACE2 transgenic mouse model

**DOI:** 10.1101/2021.04.20.440722

**Authors:** Brittany Seibert, C. Joaquín Cáceres, Stivalis Cardenas-Garcia, Silvia Carnaccini, Ginger Geiger, Daniela S. Rajao, Elizabeth Ottesen, Daniel R. Perez

**Author notes:** Address correspondence to Elizabeth Ottesen, and Daniel R. Perez.

## Abstract

Transmission of the severe acute respiratory syndrome coronavirus 2 (SARS-CoV-2), has resulted in millions of deaths and declining economies around the world. K18-hACE2 mice develop disease resembling severe SARS-CoV-2 infection in a virus dose-dependent manner. The relationship between SARS-CoV-2 and the intestinal or respiratory microbiome is not fully understood. In this context, we characterized the cecal and lung microbiome of SARS-CoV-2 challenged K18-hACE2 transgenic mice in the presence or absence of treatment with the M^pro^ inhibitor GC376. Cecum microbiome showed decreased Shannon and Inv Simpson diversity index correlating with SARS-CoV-2 infection dosage and a difference of Bray-Curtis dissimilarity distances among control and infected mice. Bacterial phyla such as Firmicutes, particularly Lachnospiraceae and Oscillospiraceae, were significantly less abundant while Verrucomicrobiota, particularly the family Akkermansiaceae, were increasingly more prevalent during peak infection in mice challenged with a high virus dose. In contrast to the cecal microbiome, the lung microbiome showed similar microbial diversity among the control, low and high challenge virus groups, independent of antiviral treatment. Bacterial phyla in the lungs such as Bacteroidota decreased while Firmicutes and Proteobacteria were significantly enriched in mice challenged with a high dose of SARS-CoV-2. In summary, we identified changes in the cecal and lung microbiome of K18-hACE2 mice with severe clinical signs of SARS-CoV-2 infection.

**IMPORTANCE:** The COVID-19 pandemic has resulted in millions of deaths. The host’s respiratory and intestinal microbiome can affect directly or indirectly the immune system during viral infections. We characterized the cecal and lung microbiome in a relevant mouse model challenged with a low and high dose of SARS-CoV-2 in the presence or absence of an antiviral M^pro^ inhibitor, GC376. Decreased microbial diversity and taxonomic abundances of the phyla Firmicutes, particularly Lachnospiraceae, correlating with infection dosage was observed in the cecum. In addition, microbes within the family Akkermansiaceae were increasingly more prevalent during peak infection, which is observed in other viral infections. The lung microbiome showed similar microbial diversity to the control, independent of antiviral treatment. Decreased Bacteroidota and increased Firmicutes and Proteobacteria were observed in the lungs in a virus dose-dependent manner. These studies add to a better understanding of the complexities associated with the intestinal microbiome during respiratory infections.

## INTRODUCTION

Throughout 2020, the World Health Organization reported ∼8 million confirmed COVID-19 cases and ∼1.8 million confirmed deaths leading to a continuous increase of cases during the early months of 2021 (1). The SARS-CoV-2 virus replicates and migrates to multiple tissues including the airways and alveolar epithelial cells in the lungs, triggering a strong immune response that may lead to exacerbation of inflammatory responses, a major complication in SARS-CoV-2 patients (2-9). While many infected patients can present as asymptomatic, others show clinical manifestations such as fever, shortness of breath, cough, headache, and occasional gastrointestinal symptoms (10-12); however, there are still several aspects of the host immune response that need to be elucidated.

The respiratory and intestinal microbiome can have direct impacts on host cells or an indirect impact on the immune system during viral infections (13, 14). Our knowledge of the microbiota’s role in essential physiological processes and disease progression has expanded greatly due to advanced sequencing technology (15), but remains poorly parameterized for many diseases. Previous studies have shown that the residential bacterial communities that reside in the respiratory tract can affect and/or be affected by respiratory viral infections, such as playing a role in the enhancement of influenza virus transmission by promoting environmental stability and infectivity (16, 17). Changes in the respiratory microbiome during influenza infection in mice showed decreased of Alphaproteobacteria and increased of Gammaproteobacteria, Actinobacteria, and facultative anaerobes like *Streptococcus* and *Staphylococcus* (16). While the potential for respiratory diseases to impact the residential microbiome is clear, many studies have also observed impacts on the intestinal microbiome during respiratory infections (18-21). Reported changes in the intestinal microbiome include enrichment of Bacteroides and Proteobacteria along with a decrease in Firmicutes during respiratory viral infections such as influenza and respiratory syncytial virus (18-21). Not only are changes observed in the intestinal bacterial communities, but it was demonstrated that TLR5 sensing of flagellated microbes in the intestine increased antibody responses post influenza vaccination and the oral administration of gut microbe *Akkermansia muciniphila* reduces weight loss and mortality during highly pathogenic influenza infection (22, 23). It has also been suggested that microbiome changes or ‘gut dysbiosis’ can lead to gut permeability resulting in secondary infections such as pneumococcal disease (20, 21). Therefore, we were interested in exploring the impacts of SARS-CoV-2 infection on the microbiome through the use of a mouse model.

The relationship between SARS-CoV-2 and the intestinal or respiratory environment is not fully understood, particularly its impact on the microbiome. Previous studies have looked at SARS-CoV-2-induced changes in the nasopharynx and fecal microbiome of humans with conflicting results (5, 24-28). Microbiome diversity and composition differences were not observed in the nasopharynx of negative and positive PCR patients in one study (28). However, another study found a decrease in the nasopharyngeal microbial diversity and differences were linked to disease severity (27). Impacts on the fecal microbiome are also expected, as numerous studies have observed viral RNA in the feces of infected individuals (9, 29), and gastrointestinal upset reported occasionally during COVID-19 infection (29). Overall, fecal microbiome studies have found a decrease in the gut microbiota diversity and abundance in SARS-CoV-2 patients compared to negative patients (5, 24, 26). Multiple bacterial genera that are associated with opportunistic pathogens such as *Streptococcocus, Rothia, Veillonella, Erysipelatoclostridium, Actinomyces, Collinsella* and *Morganella*, had increased relative abundance in fecal samples collected from SARS-CoV-2 patients compared to the controls (5, 24). Further, a recent study showed that the addition of oral bacteriotherapy treatment in human patients with SARS-CoV-2 displayed decreased mortality and reduced ICU hospitalizations (30). This suggests that understanding the host microbial changes during SARS-CoV-2 infection could help provide future treatment methods to overcome severe infections.

While nasopharynx and fecal samples can be informative, several studies in human and animal models suggest that the intestinal lumen and mucosa may be colonized by microbial communities that are different from rectal swabs or feces (31-33). However, deep respiratory and intestinal samples are more difficult to collect among human patients. Further, the human microbiome is highly variable and impacted by diverse environmental conditions (34-37), which complicates analysis of human population studies. Therefore, analyzing the respiratory and intestinal microbiome of an animal model susceptible to SARS-CoV-2 in a controlled environment that mirrors mild or severe SARS-CoV-2 infection in humans would be beneficial in understanding the relationship between SARS-CoV-2 infection and the host microbiome.

Recent reports showed that K18-hACE2 mice develop disease resembling severe SARS-CoV-2 infection in a virus dose-dependent manner, mirroring partially what is observed in humans (38-46). We aimed to use this model to understand microbiome responses to SARS-CoV-2, particularly infection or antiviral induced changes in the intestinal and lung microbiome. The studies were performed in the context of mice challenged with two different doses of the SARS-CoV-2 virus and either receiving antiviral therapy with the M^pro^ inhibitor GC-376 or vehicle for 7 days post virus challenge. We performed 16S sequencing at 2-, 5-, and 14-days post challenge (dpc) with a prototypic SARS-CoV-2 strain. The results of the intestinal microbiome show microbial differences in alpha and beta diversity measures that are SARS-CoV-2 virus dose-dependent and with little effect of GC-376 treatment on lung bacterial communities.

## RESULTS

### Clinical outcomes of K18 hACE2 transgenic mice challenged with two different doses of SARS-CoV-2 virus and samples for microbiome analyses

Taking advantage of a study evaluating antiviral activity of GC-376 against SARS-CoV-2 virus in the K18-hACE2 mouse model, we evaluated the microbiome composition at different times after SARS-CoV-2 challenge. We and others have shown that mice challenged with 10^3 TCID50/mouse of the SARS-CoV-2 virus (Low/Vehicle) presented with brief reduced activity and clinical signs leading to ∼60% survival (43). In contrast, mice challenged with 10^5 TCID50/mouse of SARS-CoV-2 (High/Vehicle) presented initially with relatively normal activity followed by rapid weight loss and substantial deterioration of clinical outcomes (43). By 6 dpc, mice in the high virus dose group showed ∼20% weight loss and all mice died or had to be euthanized by 8 dpc (43). Peak virus titers for the low and high dose groups were observed at 2 and then 5 dpc in the nasal turbinate’s and lungs (43). Antiviral GC-376 treatment resulted in milder inflammation and reduced lesions and viral loads compared to the vehicle group, although it did not improve clinical outcomes (43). We analyzed the changes in the intestinal and respiratory microbiome by collecting cecum and lungs from mice of the following groups: PBS/Vehicle, Low/Vehicle, and High/Vehicle (Fig. 1A). Because the respiratory tract is the primary site of replication for SARS-CoV-2, we also collected lung samples from the antiviral GC-376 treated group (Mock/GC-376, Low/GC-376, High/GC-376) to evaluate whether antiviral intervention would affect the residential respiratory microbiome (Fig. 1A).

**Figure 1.**
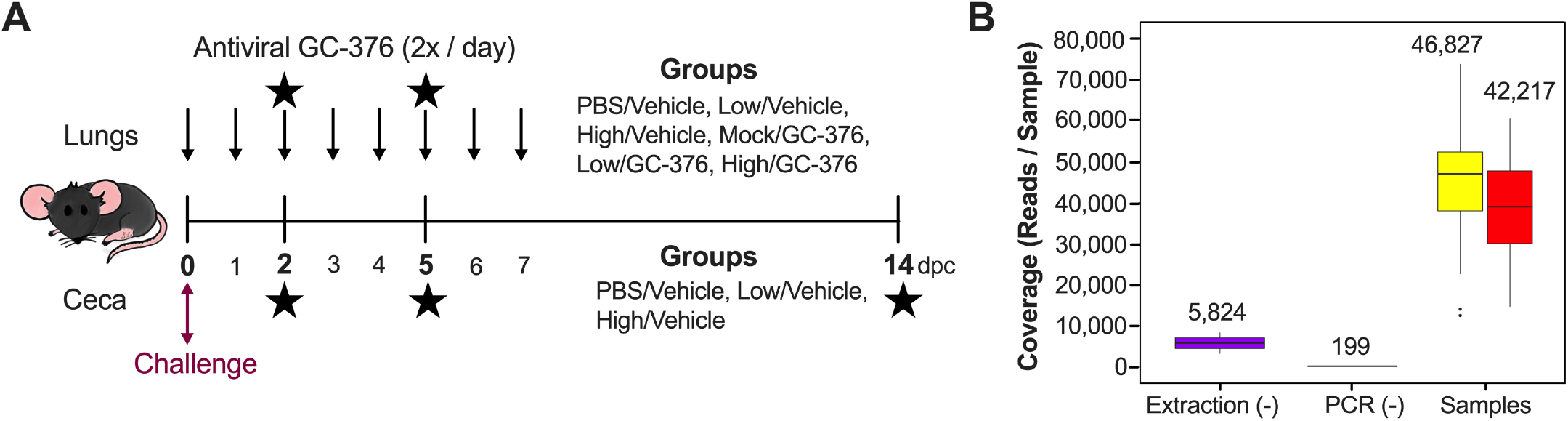
Study design and sample coverage across ceca and lung samples. **(A)** Study timeline for the mouse study. Six-week-old female mice were inoculated with PBS, low or high titers of SARS-CoV-2 virus. Three groups of mice were administered antiviral GC-376 twice per day starting 3 h after inoculation until 7 dpc (indicated by top arrows). Lung and ceca samples were collected at 2, 5, and 14 dpc (indicated by the stars). Lung samples were collected in vehicle and GC-376 groups while ceca samples were collected from the vehicle group. **(B)** Sequencing coverage of the extraction blank, PCR blank, and samples (cecum = yellow, lung = red). Coverage mean is indicated above the boxplot. Outliers are indicated by points outside of the plot. Three outliers for the ceca (above 80,000) are not shown.

We performed 16S sequencing at 2-, 5-, and 14-dpc except for lung samples in the PBS/Vehicle group due to limited DNA concentrations. Of the total ceca and lung combined 5,098,781 raw reads obtained, while 3,103,597 reads remained after dada2 trimming, filtering, merging, and chimera removal. Two lung samples, one from the Low/Vehicle at 2 dpc and one from the High/Vehicle at 5 dpc were removed from the analysis because of low coverage (<10,000 reads). One lung sample from the Mock/GC-376 at 14 dpc, considered an outlier according to Grubbs test on taxonomic abundance (p = 6.022e-07), was also removed from the analysis. Because the microbiome of the lungs can be easily contaminated, we compared the sequencing coverage of the blank extractions and negative PCR controls to samples from the lung and ceca (Fig. 1B). Blank extraction samples had an average of 5,824 reads/sample and the negative PCR controls had an average of 199 reads/sample. Meanwhile, cecum samples had an average of 46,827 with a minimum of 12,892 reads/sample and the lung samples had an average of 42,217 with a minimum of 14,808 reads/sample (Fig. 1B). Since the number of reads for the ceca and lung are notably greater than the blanks, the difference in coverage suggests that the majority of the reads in the samples are not from cross-contamination.

### Microbial diversity in the cecum of SARS-CoV-2 challenge K18 hACE2 mice

To evaluate spatial differences in microbial diversity and community structure in the cecum, we evaluated alpha diversity using count data from rarified ASVs to calculate the number of observed variants, and the Shannon and Inv Simpson indexes. There were no significant differences in ASV richness among PBS/Vehicle, Low/Vehicle and High/Vehicle when samples from each dpc were combined (Fig. 2A). Analyses of each group at each time point showed a trend towards increased number of ASVs as dpc increased; however, statistical testing was not performed because of limited sample size per time point (n= 2 or 3) (Fig. 1SA). Shannon and Inv Simpson indexes varied significantly between groups (Krustal-Wallis; p=0.015; p=0.012 respectively, Fig. 2B and 2C). The PBS/Vehicle group had the highest Shannon and Inv Simpson index, followed by Low/Vehicle, and then High/Vehicle (Fig. 2B and 2C). Pair-wise comparisons showed that PBS/Vehicle and Low/Vehicle had significantly higher Shannon and Inv Simpson diversity indexes compared to High/Vehicle (Wilcox-rank test; p = 0.015 and p=0.012; p=0.0087 and p=0.02 respectively, Fig. 2B and 2C). Shannon diversity and Inv Simpson of each group at each time point showed a similar trend among days after challenge (Fig. 1SB and 1SC). Moreover, there was no statistical difference among 2-, 5-, and 14-dpc when analyzing the virus-challenge groups (Low/Vehicle and High/Vehicle, Fig. 2S). Taken together, the alpha diversity indexes showed that the microbial diversity in the cecum of K18-hACE2 mice correlates inversely with SARS-CoV-2 virus challenge dose.

**Figure 2.**
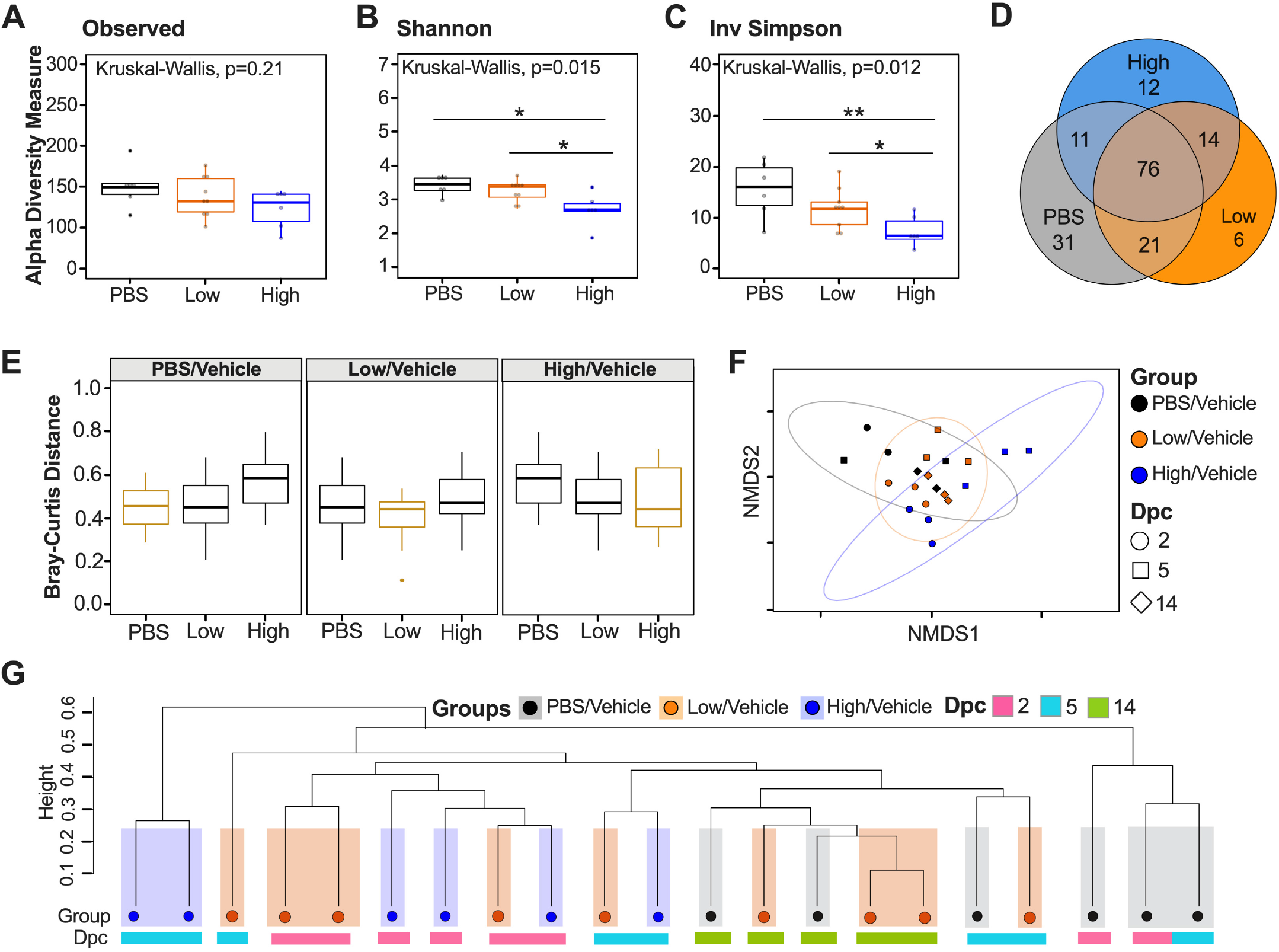
Alpha and beta diversity metrics of ceca samples. Comparison of **(A)** Observed ASVs, **(B)** Shannon diversity index and **(C)** Inv Simpson of different groups (PBS/Vehicle: Black, Low/Vehicle: Orange, High/Vehicle: Blue) containing all dpc from rarified ASV count table. **(D)** Venn diagram of rarified ASV counts comparing the three different groups. **(E)** Comparison of weighted Bray-Curtis dissimilarity distances within each group and across different groups. Gold boxes represent within-group variation while black boxes represent the between-group variation. **(F)** NMDS plot of weighted Bray-Curtis dissimilarity distance. Days post challenge are indicated by the shape and groups are indicated by color. Ellipses were constructed using a multivariate t-distribution. **(G)** Dendrogram showing the relationship of different groups and dpc using Bray-Curtis dissimilarity distance. Hierarchical cluster analysis was performed using hclust with agglomeration method average. Shaded colors and circles correspond to the different groups as described previously. Colored bars below the circles represent the different dpc (pink= 2 dpc, light blue = 5 dpc, green = 14 dpc). All statistical tests were performed using Kruskal-Wallis or Wilcox-rank test for pair-wise comparisons using * = p<0.05 ; ** = p<0.005; *** = p<0.0005; **** = p<0.00005.

In order to assess the relationship between microbial community structure and SARS-CoV-2 challenge during the course of infection, we analyzed the number of shared ASVs. The three groups shared 76 ASVs after the count data was rarified with a detection limit of 0.001 in at least 90% of the samples (Fig 2D). Using the same criteria, PBS/Vehicle had 31 unique ASVs while Low/Vehicle and High/Vehicle had 6 and 12 unique ASVs, respectively (Fig 2D). The Low/Vehicle group shared 21 ASVs with PBS/Vehicle and 11 ASVs with High/Vehicle, suggesting a more similar microbial composition between Low/Vehicle and PBS/Vehicle (Fig. 2D). Next, we quantified changes of the cecum microbiome composition among different SARS-CoV-2 infected groups by comparing weighted dissimilarity distances (Bray-Curtis) within and across groups (Fig. 2E). Overall, High/Vehicle showed greater dissimilarity to PBS/Vehicle than Low/Vehicle in all group comparisons (Fig. 2E). A non-metric Multi-dimensional Scaling (NMDS) plot of the Bray-Curtis dissimilarity distance was used to assess the relationship between microbial community structure and SARS-CoV-2 challenge during the course of infection. The NMDS showed a spread of samples; however, the PBS/Vehicle showed more overlap with the Low/Vehicle compared to the High/Vehicle, which was further supported by a PERMANOVA analysis analyzing difference of groups (p=0.001) (Fig. 2F). We further investigated the relationship of the three groups across dpc by producing a hierarchical cluster analysis using the Bray-Curtis dissimilarity distances (Fig. 2G). While the samples did not cluster exclusively by treatment, in all cases the High/Vehicle clustered with samples within their group or with Low/Vehicle treatment samples to the exclusion of PBS/Vehicle, and similarly PBS/Vehicle clustered with their own group or Low/Vehicle to the exclusion of High/Vehicle (Fig. 2G). Collectively, the beta diversity metrics suggest that a higher dosage of SARS-CoV-2 virus infection has a larger effect on microbial diversity and community structure compared to a low virus dose infection.

Since the diversity metrics suggested a difference among the low and high virus dose infected mice, we investigated those differences further by analyzing the relative abundance of the microbial communities at the phylum and family levels. The most abundant phyla were Bacteroidota, Firmicutes, Proteobacteria, and Verrumicrobiota. The PBS/Vehicle group had significantly greater relative abundance of Firmicutes compared to Low/Vehicle and High/Vehicle (Wilcox-rank test; p=0.025; p=0.0087 respectively) (Fig. 3A). The Low/Vehicle group had significantly more abundant Proteobacteria compared to High/Vehicle (Wilcox-rank test; p=0.0008) (Fig. 3A). While not statistically significant, High/Vehicle had notably higher abundance of Verrucomicrobiota compared to the other two groups. Next, we looked at the relationship between the two predominant phyla to calculate the Firmicutes/Bacteroidota (F/B) ratio. The PBS/Vehicle group had the highest F/B ratio, followed by Low/Vehicle, and then High/Vehicle (Fig. 3B). Pair-wise comparisons showed that PBS/Vehicle had significantly higher F/B ratio compared to High/Vehicle (Wilcox-rank test; p = 0.02, Fig. 3B). When analyzing taxonomic diversity at the family level, the most abundant families were Muribaculaceae, Lachnospiraceae, and Akkermansiaceae (Fig. 3B left). Muribaculaceae was similar among groups when all sample time points were combined, consistent with its parent phyla, Bacteroidota (Fig. 3A). The PBS/Vehicle and Low/Vehicle had significantly higher abundances of Lachnospiraceae compared to the High/Vehicle (Wilcox-rank test; p=0.05, p=0.041 respectively, Fig. 3B left). Another family in the phyla Firmicutes, Oscillospiraceae, had significantly increased abundance in the PBS/Vehicle group compared to the High/Vehicle (Wilcox-rank test; p=0.0087, Fig. 3B right). While not significant, the family driving the increase shift of Verrumicrobiota in the High/Vehicle was Akkermansiaceae (Fig. 3B left). This family was notably enriched at 5 dpc in 2 out of 3 samples of the High/Vehicle (Fig. 3D). Taken together, the taxonomic relative abundances displayed distinct changes at the phylum and family level for the high dose infected group while the control and low dose groups were more similar, particularly in families within the phylas’ Bacteroidota, Firmicutes, Proteobacteria, and Verrumicrobiota.

**Figure 3.**
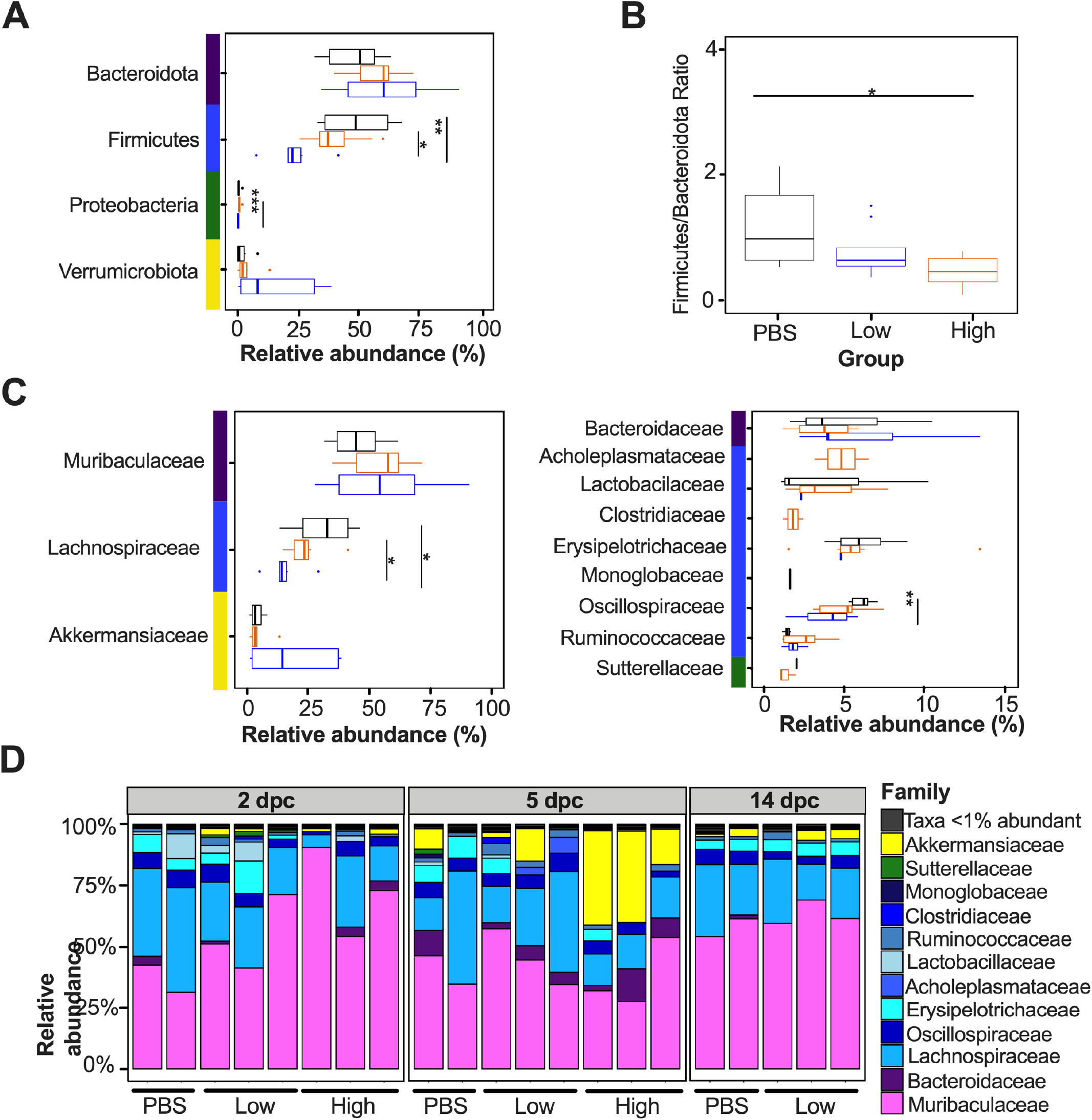
Relative abundance (%) of microbial communities in the ceca at the phylum and family level. **(A)** Relative abundances (%) of the most abundant phyla were compared via box plots. Each box represents the interquartile range (first and third quartiles) of taxa abundance, and the line corresponds to the median abundance. Vertical lines represent variation in abundance and the circles represent outliers. Corresponding phyla are noted by the colored bar to the left of the graphs (purple = Bacteroidota, blue = Firmicutes, green = Proteobacteria, yellow = Verrumicrobiota). **(B)** Firmicutes/Bacteroidota ratio was calculated and graphed to analyze differences among different groups. **(C)** Relative abundances (%) of the most abundant families were compared via box plots. Corresponding phyla are noted by the colored bar to the left of the graphs following legend in A. **(D)** Relative abundances (%) of individuals were calculated by agglomerating at the family level and then transformed into relative abundances. Taxa that had less than 1% abundance was grouped together. Groups are indicated by the bars at the bottom of the graph and dpc at the top of the graph. All statistical tests were performed using Wilcox-rank test using * = p<0.05 ; ** = p<0.005; *** = p<0.0005; **** = p<0.00005.

### Microbial diversity in the lungs of SARS-CoV-2 challenge K18 hACE2 mice

Since we observed intestinal dysbiosis in SARS-CoV-2 infected mice, we subsequently analyzed the changes of the antiviral M^pro^ inhibitor GC-376 and infection in the lung microbiome. The effect of GC-376 was analyzed from homogenized lung tissue samples and we specifically investigated how antiviral treatment may have affected microbiome changes in the lung during SARS-CoV-2 infection. Due to low DNA concentrations and <10,000 reads/sample, we did not include the six lung samples from the PBS/Vehicle group in our analyses. Therefore, analysis of the microbiome was only assessed in lung samples from Low/Vehicle, High/Vehicle, Low/GC-376, and High/GC-376. The low virus challenge dose groups showed no significant differences among number of observed ASVs, Shannon or Inv Simpson diversity indexes between GC-376-treated and untreated mice when samples from all time points were taken into account (Fig. 4A, B, and C). Bray-Curtis dissimilarity distances similarly did not show treatment-specific differences (Fig. 4D; PERMANOVA p=0.88). Results from the two high virus challenge dose groups showed that High/Vehicle had significantly higher combined number of ASVs compared to High/GC-376 (Fig. 4E). However, the Shannon and Inv Simpson diversity indexes were similar (Fig. 4F and G), and the Bray-Curtis dissimilarity did not cluster by treatment group or show treatment-specific differences (Fig. 4H; PERMANOVA p=0.72). Collectively, the results showed that the lung microbial communities during infection with SARS-CoV-2 were unaffected by the antiviral treatment at low or high virus challenge doses.

**Figure 4.**
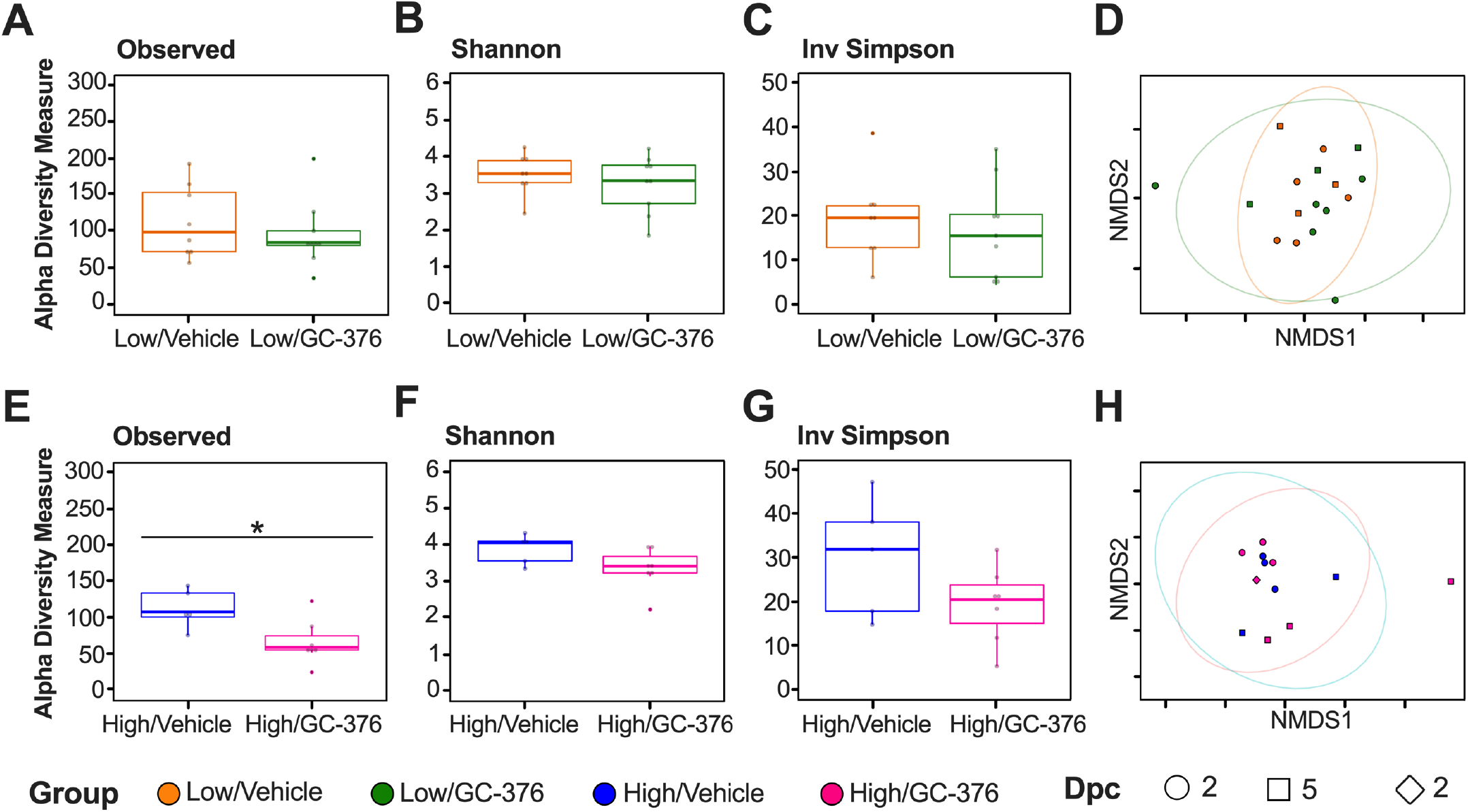
Alpha and Beta Diversity metrics of lung samples without / with antiviral GC-376. Comparison of **(A)** Observed ASVs, **(B)** Shannon diversity index and **(C)** Inv Simpson of the low dose groups without (Low/Vehicle: Orange) and with antiviral (Low/GC-376: Green) containing all dpc from rarified ASV count table. **(D)** NMDS plot of weighted Bray-Curtis dissimilarity distance of the rarified ASV count table of the low dose groups. Days post challenge are indicated by the shape and groups by color. Ellipses were constructed using a multivariate t-distribution. Comparison of **(E)** Observed ASVs, **(F)** Shannon diversity index and **(G)** Inv Simpson of the high dose groups without (High/Vehicle: Blue) and with antiviral (High/GC-376: Pink). **(H)** NMDS plot of weighted Bray-Curtis dissimilarity distance of the rarified ASV count table of the high dose groups. Days post challenge are indicated by the shape and groups by color. Ellipses were constructed using a multivariate t-distribution. All statistical tests were performed using Kruskal-Wallis or Wilcox-rank test for pair-wise comparisons using * = p<0.05 ; ** = p<0.005; *** = p<0.0005; **** = p<0.00005.

Because the PBS/Vehicle treatment did not yield high-quality data and similar lung microbial communities were observed regardless of whether antiviral treatment took place, alpha and beta diversity metrics for the lung microbiome were compared for the GC-376 treated groups (Mock/GC-376, Low/GC-376 and High/GC-376) to examine the impact of infection on the lung microbiome. The number of observed ASVs (Krustal-Wallis; p=0.041; Fig. 5A) varied significantly among groups. The Mock/GC-376 group had the highest number of ASVs, followed by Low/GC-376 and then High/GC-376; the Mock/GC-376 group had significantly greater number of ASVs compared to High/GC-376 including all time points (Wilcox-rank test; p=0.021) (Fig. 5A). Analyses at different dpc revealed similar number of ASVs regardless of time point for the Mock/GC-376, whereas virus challenge groups had decreased number of ASVs as the infection progressed (Fig. 3S). In contrast to the ceca, Shannon and Inv Simpson indexes for the lung microbiome of the GC-376-treated groups were not significantly different when comparing across viral doses or over the course of infection (Fig. 5B and 5C; Fig. 3S and 4S).

**Figure 5.**
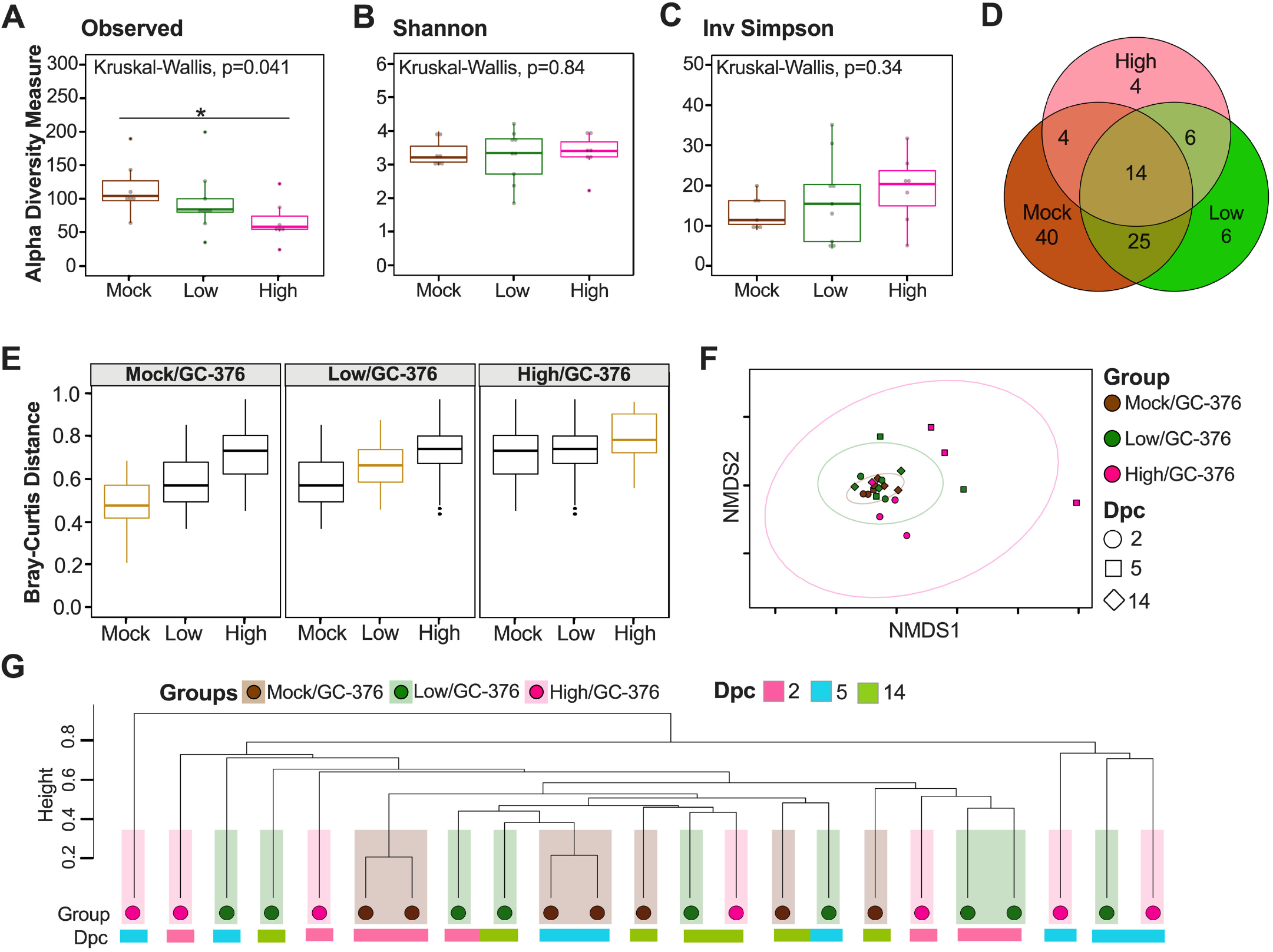
Alpha and beta diversity metrics of lung samples. Comparison of **(A)** Observed ASVs, **(B)** Shannon diversity index and **(C)** Inverse Simpson of different groups (Mock/GC-376: Brown, Low/GC-376: Green, High/GC-376: Pink) containing all dpc from rarified ASV count table. **(D)** Venn diagram of rarified counts comparing the three different groups. **(E)** Comparison of Bray-Curtis dissimilarity distances within each group and across different groups. Gold boxes represent within variation while black boxes represent other groups. **(F)** NMDS plot of weighted Bray-Curtis dissimilarity distance of the rarified ASV count table. Days post challenge are indicated by the shape and group by color. Ellipses were constructed using a multivariate t-distribution. **(G)** Dendrogram showing the relationship of different groups and dpc using Bray-Curtis dissimilarity distance. Hierarchical cluster analysis was performed using hclust with agglomeration method average. Shaded colors and circles correspond to the different groups described previously. Colored bars below the circles represent the different dpc (pink= 2 dpc, light blue = 5 dpc, green = 14 dpc). All statistical tests were performed using Kruskal-Wallis or Wilcox-rank test for pair-wise comparisons using * = p<0.05 ; ** = p<0.005; *** = p<0.0005; **** = p<0.00005.

Since the alpha diversity analysis suggest limited to no differences among SARS-CoV-2 infected mice, we next analyzed the number of shared ASVs among the different groups (Fig. 5D). The three GC-376-treated groups shared 14 ASVs (rarified count data, detection limit of 0.001 in at least 90% of the samples, Fig. 5D). Following the same criteria, the Mock/GC-376 group had 40 unique ASVs while the Low/GC-376 and the High/GC-376 had 6 and 4 unique ASVs, respectively (Fig. 5D). The Low/GC-376 shared 25 ASVs with the Mock/GC-376 and 6 ASVs with the High/GC3-76 (Fig. 5D). In order to further understand the differences among groups, we quantified the change of the lung microbiome composition among different groups by comparing Bray-Curtis distances within and across groups (Fig. 5E). The results suggest that the High/GC-376 were most dissimilar to the others, while the Low/GC-376 group and the PBS/GC-376 group were more similar (Fig. 5E). In addition, the High/GC-376 showed greater within-group variation than Low/GC-376 and Mock/GC-376 (Fig. 5E). Next, a NMDS plot of the Bray-Curtis dissimilarity distance was used to assess the relationship among the lung microbial community and SARS-CoV-2 challenge during the course of infection (Fig. 5F). Bray-Curtis dissimilarity NMDS showed tight grouping of samples with outliers that belonged to the High/GC-376 group at 5 dpc, which was supported by a PERMANOVA analysis analyzing difference of groups (p=0.01) (Fig. 5F). Further, we investigated the relationship of the three groups across dpc by producing a hierarchical cluster analysis using the Bray-Curtis dissimilarity distances (Fig. 5G). The lung microbiota showed less clustering by treatment than the ceca (Fig. 5G). Altogether, the results suggest that there are dose-dependent changes in microbial community composition following SARS-CoV-2 infection, although the results are less clear-cut than observed in the cecal microbiome.

Considering the diversity metrics suggested a limited difference in infected and control mice, we further analyzed the relative abundance of the microbial communities at the phylum and family levels. The most abundant phyla within the lungs were Bacteroidota, Firmicutes, Proteobacteria, Actinobacteriota and Verrumicrobiota. In contrast to the ceca, Bacteroidota were suppressed in GC-376-treated mice exposed to low and high dose virus, with the Mock/GC-376 exhibiting significantly higher abundance of Bacteroidota compared to High/GC-376 (p=0.017) (Fig 6A). The High/GC-376 group had significantly higher abundance of Firmicutes than Mock/GC-376 (Wilcox-rank test; p=0.038) (Fig. 6A). The Low/GC-376 and High/GC-376 had significantly more abundant Proteobacteria compared to Mock/GC-376 (Wilcox-rank test; p=0.025; p=0.0087 respectively, Fig. 6A). Similar abundances were observed for Actinobacteriota and Verrucomicrobiota across all groups (Fig. 6A). Despite having similar if not lower ASV-level diversity, lung samples showed higher family-level diversity than the cecum.

**Figure 6.**
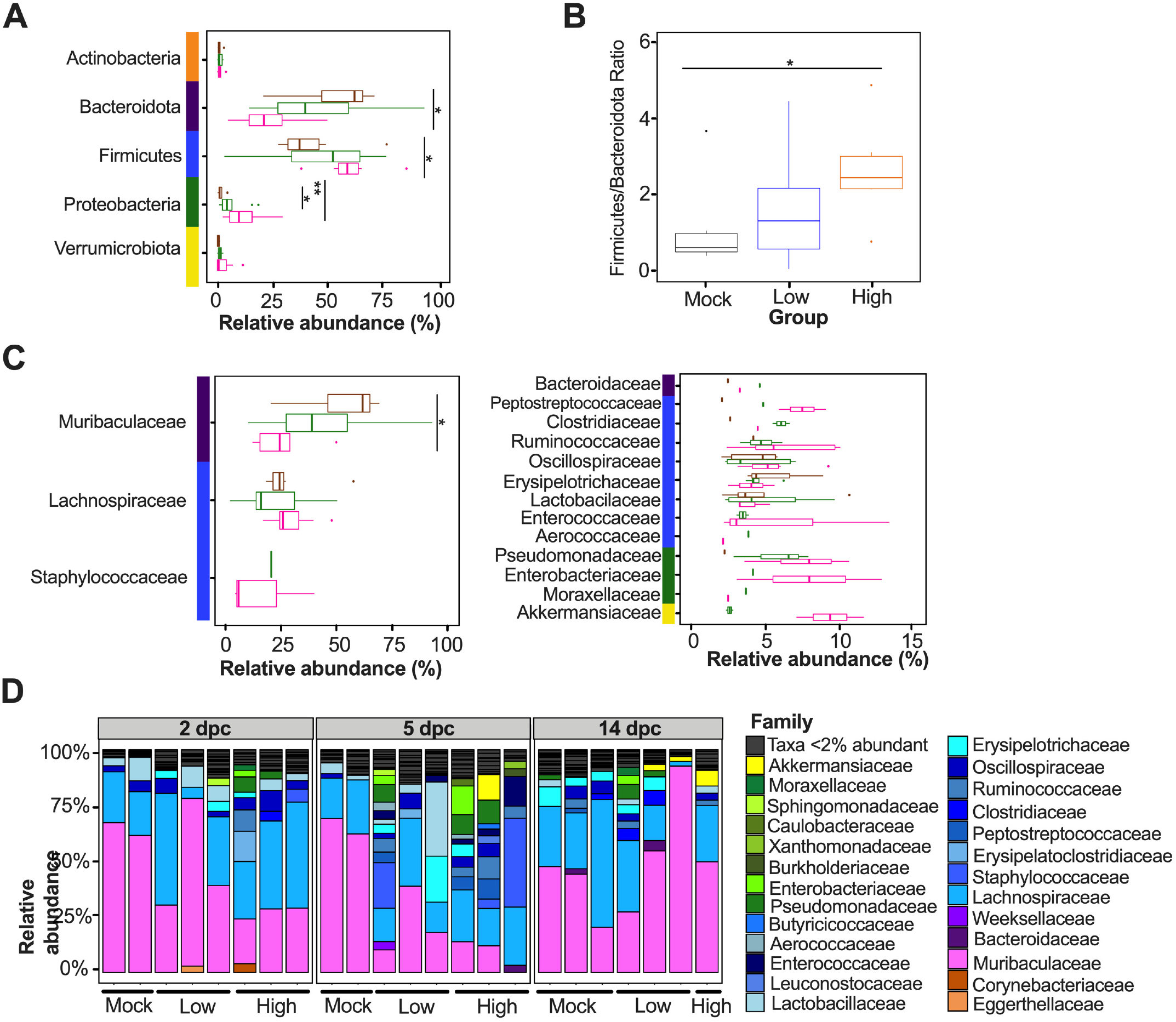
Relative abundance (%) of microbial communities in the lung at the phylum and family level. **(A)** Relative abundances (%) of the most abundant phyla were compared via box plots. Each box represents the interquartile range (first and third quartiles) of taxa abundance, and the line corresponds to the median abundance. Vertical lines represent variation in abundance and the circles represent outliers. Corresponding phyla are noted by the colored bar to the left of the graphs (purple = Bacteroidota, blue = Firmicutes, green = Proteobacteria, yellow = Verrumicrobiota). **(B)** Firmicutes/Bacteroidota ratio was calculated and graphed to analyze differences among groups. **(C)** Relative abundances (%) of the most abundant families were compared via box plots. Corresponding phyla are noted by the colored bar to the left of the graphs following legend in A. **(D)** Relative abundances (%) of individuals were calculated by agglomerating at the family level and then transformed into relative abundances. Taxa that had less than 2% abundance was grouped together. Groups are indicated by the bars at the bottom of the graph and dpc at the top of the graph. All statistical tests were performed using Wilcox-rank test for pair-wise comparisons using * = p<0.05 ; ** = p<0.005; *** = p<0.0005; **** = p<0.00005.

Following, we looked at the relationship among Firmicutes and Bacteroidota by analyzing the F/B ratio. Contrary to the ceca, the Mock/GC-376 group had the lowest F/B ratio, followed by Low/GC-376, and then High/GC-376 (Fig. 6B). Pair-wise comparisons showed that Mock/GC-376 had significantly lower F/B ratio compared to the High/Vehicle (Wilcox-rank test; p = 0.02, Fig. 6B). When analyzing taxonomy at the family level, the most abundant families in the lungs were Muribaculaceae, Lachnospiraceae, and Staphylococcaceae (Fig. 6B left and 6C). Across all dpc, Mock/GC-376 had significantly higher abundance of Muribaculaceae compared to High/GC-376, like its parent phyla, Bacteroidota (Wilcox-rank test; p=0.03, Fig. 6B and 6C left). All groups in the lungs had similar abundances of Lachnospiraceae and Staphylococcaceae (Fig. 6C left and 6D). Collectively, the taxonomic relative abundances displayed distinct changes at the phylum and family level in mice challenged with low and high challenge doses of the SARS-CoV-2 virus and treated with GC-376, particularly in families within the phylas’ Bacteroidota, Firmicutes, and Proteobacteria.

## DISCUSSION

We analyzed the cecum and lung microbiome changes that occur in K18-hACE2 mice upon challenge with two different doses of a prototypical SARS-CoV-2 virus. Some limitations of this study must be noted. While the environment was stable and controlled, the sample size for each group at each time point was small and the potential contribution of cage effect on the microbiome was not analyzed. The high mortality observed in the high virus dose groups only allowed for collection of two time points (2 and 5 dpc, only one sample was collected at 14 dpc from the only survivor in the High/GC-376 group). As indicated above, ceca samples were only collected for mice not treated with GC-376. In contrast, lung samples from mock inoculation/vehicle treated control mice did not yield sufficient amplifiable microbial DNA for sequencing, so the comparison focused on SARS-CoV-2 dose-dependent responses in GC-376 treated mice.

The microbiome of the cecum showed significant decreases in Shannon and Inv Simpson index comparing the control to the low-dose and high-dose infected groups (Fig. 2B and 2C). The low virus dose group shared a higher number of ASVs with the control group compared to the high virus dose group (Fig. 2D). These observations suggest a virus dose-dependent effect of the ceca microbial diversity in mice infected with SARS-CoV-2. While preparing the manuscript, a report was published that analyzed the small intestine microbiome of hACE2 mice among unvaccinated and vaccinated mice challenged with a high dose of SARS-CoV-2 (47). While we compare control mice to low and high doses of non-vaccinated mice challenged with SARS-CoV-2 in this study, findings were similar since a decrease in alpha diversity in the unvaccinated mice was reported, which was consistent to results obtained from human fecal samples (5, 47). From the beta diversity analysis, the weighted Bray-Curtis NMDS showed that samples from the high virus dose group were outliers compared to the rest of the samples, consistent with exacerbation of clinical signs and following peak virus replication in this group (40, 43, 44, 48). Further analysis of the Bray-Curtis distances of cecal samples showed that the low dose group was more similar to the control group compared to the high dose group (Fig. 2F and G) suggesting that microbial change is virus dose-dependent. A similar relationship was previously observed among unvaccinated and vaccinated SARS-CoV-2 infected mice with a high dose (47). Analyzing the commensal microbiome in other diseases have shown that the microbiota can both regulate and be regulated by viral pathogens and facilitate stimulatory or suppressive effects on the host immune response (49). It is possible that the distinct clustering and change observed at 5 dpc in the cecal microbiome of mice infected with a high virus dose could be caused by a hyperactive host innate immune response and/or SARS-CoV-2 virus replication. It is also possible that cecal microbiome changes could also contribute to the rapid increase of disease severity. Viral pathogens that infect or replicate in mucosal tissues most likely encounter commensal microbiota inhabiting the mucosal surfaces (50). Therefore, the intestinal microbiota can either promote viral infections such as poliovirus, reovirus and certain retroviruses, or it can have a protective role such as influenza and rotavirus (51-54).

The most distinct differences in taxonomic relative abundance within the ceca of infected mice is the overall lower abundance of Firmicutes, particularly the families Lachnospiraceae and Oscillospiraceae, increased abundance of Proteobacteria in the low virus dose group, and the increase of Verrucomicrobiota, particularly the family Akkermansiaceae, in the high virus dose group at 5 dpc (Fig. 3). In addition, a significant difference among the F/B ratio was observed among the control and high dose group (Fig 3B). Previously, F/B ratio has been associated with maintaining homeostasis and changes could be indicative of dysbiosis (55). The results in this report showed a decrease in the F/B ratio within the ceca, which was also observed in patients with inflammatory bowel disease and in mice infected with RSV (20, 55). Multiple studies have reported a decrease in Firmicutes during respiratory viral infections in the intestinal microbiome, particularly in influenza infection (5, 18, 20). Interestingly, Firmicutes, particularly Lachnospiraceae, was not significantly decreased in SARS-CoV-2 stool samples compared to the controls in humans (5). Members of the Lachnospiraceae family are anaerobic, fermentative bacteria that hydrolyze starches, sugars and other short chain fatty acids (SCFAs) (56).

Previous reports have shown that SCFAs are important for the maintenance of colonic epithelial cells, directly interacts with the host immune response and promote bactericidal activity of alveolar macrophages during influenza infection (21, 56). As observed in this study, the decrease in Firmicutes, particularly Lachnospiraceae, correlates with virus challenge dose (Fig. 3). Therefore, variations in analyses of human samples could be dependent on sample type and viral load during infection as shown within this study. In addition, previous studies have shown increased Proteobacteria during influenza infection in mice, similar to the results in this study (19, 57). The increase in Proteobacteria has been hypothesized to be mediated by type 1 interferons, which has been shown to be impaired in severe SARS-CoV-2 cases but not in patients with mild-moderate outcomes (58, 59). The decrease of type 1 interferon in severe patients could potentially correlate with the significant increase of Proteobacteria in the low virus dose compared to the high virus dose found within this study; however, further research is needed to investigate this relationship. Finally, increases in abundance of the family Akkermansiaceae is of particular interest. The family Akkermansiacea was classified further into one genus *Akkermansia* (60). A previous report showed that hACE2 mice that were not vaccinated and challenged with SARS-CoV-2 had significantly increased *Akkermansia* compared to vaccinated challenged mice, similar to what is observed in this study among controls and the high challenge dose group (47). One of the primary *Akkermansia* species in the cecum of mice, *Akkermansia muciniphila*, a gram-negative, obligate anaerobe, has been shown to alter mucosal gene expression towards increased expression of genes involved in the immune response, particularly genes involved in antigen presentation of leukocytes (61). As expected, mice from the high virus dose group had increased staining of CD3^+^ and Iba-1^+^ cells, markers for cellular infiltration, and more pronounced neutrophilic inflammation compared to the low virus dose and control (43). While 2 of the 3 high virus dose mice at 5 dpc were outliers compared to the other samples, one mouse was closer in proximity to the low virus dose and control group in the Bray-Curtis NMDs and had a more similar taxonomic composition to these groups (Fig. 2F and G and Fig. 3C). Looking closer at the taxonomic differences, the sample from this mouse also had lower relative abundance of *Akkermansia* compared to the other two mice at 5 dpc (Fig. 3C). While challenged with the same dose, this mouse showed less clinical signs throughout infection, thus, suggesting that the decreased abundance of *Akkermansia* in this mouse could be correlative to disease severity. A recent report showed that while the abundance of *Akkermansia* positively correlated with influenza H7N9 infection in mice; however, oral administration of *A. muciniphila* significantly reduced weight loss, mortality, and viral titers (22). Therefore, further research is needed to better understand the role of *Akkermansia* in severe SARS-CoV-2 infection. In conclusion, the cecal data suggests that the distinct changes of the ceca microbiome could be virus dose-dependent, and specific taxa could play a role in the modulation of the immune response potentially leading to multisystemic inflammatory syndrome, a major complication of SARS-CoV-2 infection (2-9).

While the intestinal microbiome has been the center for previous microbiome research; recently, multiple groups have analyzed the microbiome composition of the upper and lower respiratory tract (16, 18, 62, 63). In particular, the lung microbiota is understood to provide resistance to the colonization of respiratory pathogens and immune tolerance (63). To our knowledge, no studies have analyzed the effects of an antiviral on the lung microbiome. We found no significant differences among groups of mice challenged with SARS-CoV-2 that were either treated or not treated with the M^pro^ inhibitor GC-376 (Fig. 4). Since GC-376 had limited effect on the clinical outcome of SARS-CoV-2 in mice, the limited differences in the microbial composition in the lungs between treated and non-treated mice is not entirely surprising. Among the samples obtained from GC-376-treated mice and in contrast to samples from ceca, differences in alpha diversity indexes between control, low-dose, and high-dose infected mice were not observed besides decreased number of ASVs within the lungs (Fig. 5A). Similar to results within this study, a previous report also showed no significant differences in Shannon diversity within the lung microbiome of mice infected with influenza and the nasopharynx of negative and positive SARS-CoV-2 PCR patients (18, 28). Regarding beta diversity analysis, the weighted Bray-Curtis NMDS did not show separation of clusters by treatment, but rather a single overlapping cluster with outliers that primarily belong to the high virus group (Fig. 5F). In contrast, a similar analysis using influenza detected no significant changes in beta diversity of the lower respiratory tract throughout infection (18). Analysis of Bray-Curtis dissimilarity indicated that the low-dose infected mice were more similar to the control group compared to the high-dose infected mice (Fig. 6F and G), and that within-group variance increased in an infectious-dose dependent manner, consistent with the Anna-Karenina model of disease-induced dysbiosis (64, 65). Comprehensively, the beta diversity analysis suggests that there are limited lung microbial composition changes, dissimilar to the ceca.

The most distinct differences in taxonomic relative abundance within the lung of infected mice is the overall lower abundance of Bacteroidota, higher abundance of Firmicutes, and higher abundance of Proteobacteria in the low and high virus dose groups compared to the mock control (Fig 6), which is consistent with previous reports (66, 67). Firmicutes and Proteobacteria were enriched in the high and low virus dose compared to the mock control, consistent with previous reports of patients infected with influenza (68). A significant difference among the F/B ratio was observed among the control and high dose group suggesting dysbiosis in the lung microbiome post SARS-CoV-2 infection with a high dose. Similar results were observed in patients that underwent lung transplants but to our knowledge, this has not been thoroughly examined in respiratory viral infections (69).

Future studies are needed with larger group sizes, cage effect compensation and analysis of different sections of the intestinal tract (duodenum, jejunum, and ileum) and respiratory tract (lower and upper sections) to better understand the role of microbiome changes during SARS-CoV-2 infection. However, the proof-of-principle approach of this report identified significant changes in the cecal and lung microbiome of K18-hACE2 mice, particularly those challenged with a high dose of the SARS-CoV-2 virus that warrants for more in-depth studies.

## MATERIALS AND METHODS

### Ethics statement

Animal studies were approved by the Institutional Animal Care and Use Committee (IACUC) of the University of Georgia (Protocol A2019-03-032-Y1-A3) and performed following the IACUC Guidebook of the Office of Laboratory Animal Welfare and PHS policy on Humane Care and of Use of Laboratory Animals. Animals were humanely euthanized following guidelines by the American Veterinary Medical Association (AVMA). Studies were performed in an animal Biosafety level 3 containment facility at the Animal Health Research Center (AHRC) at the University of Georgia.

### Cells and Virus

The SARS-CoV-2 (Isolate USA-WA1/2020) isolate, kindly provided by Dr. S. Mark Tompkins, Department of Infectious Diseases, University of Georgia, was used for virus challenge in the animal studies. Virus propagation and titration is explained in detail in Caceres et al. 2021 (43). Briefly, the virus was grown in Vero E6 Pasteur cells provided by Maria Pinto (Center for Virus research, University of Glasgow, Scotland, UK), and maintained in Dulbecco’s Modified Eagles Medium (DMEM, Sigma-Aldrich, St Louis, MO) containing 10% fetal bovine serum (FBS, Sigma-Aldrich, St Louis, MO), 1% antibiotic/antimycotic (AB, Sigma-Aldrich, St Louis, MO) and 1% L-Glutamine (Sigma-Aldrich, St Louis, MO). Cells were cultured at 37ºC under 5% CO_2_ for 96 h. Virus stocks were titrated by tissue culture infectious dose 50 (TCID_50_) and virus titers were established by the Reed and Muench method (70).

### Mouse experiments

Female K18-hACE2 mice (6 weeks old) were randomly distributed into six groups (n=6/group for controls and n=9/group for challenged), anesthetized and challenged intranasally with 50 µL of phosphate buffer saline (PBS), 1×10^3^ TCID_50_/mouse (Low virus dose) or 1×10^5^ TCID_50_/mouse (High virus dose). At 3 h post-challenge, GC-376 (20mg/kg/dose, 40 mg/kg daily), kindly provided by Dr. Jun Wang, (Department of Pharmacology and Toxicology, University of Arizona), or vehicle (H_2_O) was administered to each mouse through intraperitoneal injection (i.p.) twice per day and continued for 7 days (Fig. 1A). Mice were monitored twice a day for clinical signs of disease post challenge. Mice were humanely euthanized if they lost ≥25% of their initial body weight (a score of 3 on a 3-point scale of disease severity). At 2- and 5-dpc, a subset of mice was humanely euthanized, n=2/time point from PBS/Vehicle and Mock/GC-376 and n=3/time point from low (Low/Vehicle and Low/GC-376) and high (High/Vehicle and High/GC-376) dose. Ceca and lungs from each mouse were collected and stored at -80ºC until further analysis. At 14 dpc, the same procedure was performed with all of the remaining animals (PBS/Vehicle, Mock/GC-376, Low/Vehicle, Low/GC-376, and High/GC-376) (Fig. 1A).

### Tissue Sample Preparation

Tissue homogenates were generated using the Tissue Lyzer II (Qiagen, Gaithersburg, MD). In summary, 500 µL of PBS-AB was added to each sample (Lungs = 0.01 – 0.04g; Cecum = 0.3 -0.5g) along with Tungsten carbide 3mm beads (Qiagen, Gaithersburg, MD). Samples were homogenized at a speed of 10 Hz for 10 min. Homogenized tissue was stored at -80 until further analysis.

### DNA extraction, amplicon library preparation and sequencing

DNA was extracted from the tissue homogenates using MoBio Power Soil Kit (Qiagen, Gaithersburg, MD) with minor changes following the Earth Microbiome Protocol as follows: additional incubation at 65°C for 10 min after the addition of solution C1, beads were shaken at 20 Hz for 20 min instead of 10 min, and samples were incubated at 4°C for 10 min instead of 5 min and then stored at -80°C until use. Following extraction, the microbial 16S rRNA gene was amplified using Phusion Hot Start 2 DNA polymerase (Thermo Fisher, Waltham, MA) and V4 hypervariable region of the 16S rRNA gene primers 515F (5’-GT GCCAGCMGCCGCGGTAA -3’) and 806R (5’-GGACTACHVGGGTWTCTAAT -3’) in 20 µL PCR reactions (8.9 µL of Molecular-grade water, 4 µL of 5X HF Buffer, 0.4 µL of 10mM dNTPs, 1.25 µL of 10uM 515-F, 1.25 µL of 10uM 806R, 4 µL of DNA, 0.2 µL of polymerase) under the following conditions: 98°C (30 s), followed by 25 cycles of 98°C (10 s), 52°C (30 s), 72°C (30 s), a final elongation step at 72°C (5 min), and held at 4°C. The PCR reactions were performed in duplicate and products were visualized on a 1% agarose gel. Duplicate PCR products of the same sample were pooled in equal volumes and cleaned by 0.45x of Agencourt AMPure XP Magnetic Beads (Beckman Coulter, Pasadena, CA) according to manufacturer’s protocol, and eluted in molecular biology grade water (Genesee Scientific, San Diego, CA). Amplicon concentration was measured using the Qubit dsDNA HS Assay kit (ThermoFisher, Waltham, MA) on the Qubit 3.0 fluorometer (ThermoFisher, Waltham, MA). DNA concentrations were normalized to 1.0 ng/µL. Subsequently, amplified DNA was used in a secondary amplification/dual barcode annealing reaction. Forward and reverse dual barcode primers (primers and barcodes with different reference indices) were designed based upon primers generated by Caporaso et al. (71). Secondary amplification reactions were performed using NEBNext High-Fidelity 2X PCR Master Mix (NEB) in 50 µL reaction (26 µL of 2X Mix, 20.5 µL of water, 1uL of barcoded forward and reverse primers (10uM), 1 µL of DNA) under the following conditions: 98 °C (30 s), followed by four cycles of 98 °C (10 s), 52 °C (10 s), 72 °C (10 s), followed by six cycles of 98 °C (10 s), 72 °C (1 min), followed by a final extension of 72°C (2 min) and then held at 4°C. Samples were subsequently cleaned by 0.45x of Agencourt AMPure XP Magnetic Beads according to manufacturer’s protocol and eluted in molecular biology grade water. Fragment size distribution was analyzed on a subset of samples using the Agilent Bioanalyzer 2100 DNA-HS assay (Agilent, Santa Clara, CA, USA). Sample libraries were then normalized and pooled to a concentration of 2 or 0.5 nM based on a predicted total product size of ∼ 420 bp using the Qubit dsDNA HS Assay kit on the Qubit 3.0 fluorometer. The loading concentration of the pooled libraries was 10 pM. Libraries were sequenced using Illumina MiSeq V2 chemistry 2×250 (Illumina, San Diego, CA) paired end. Negative controls including an extraction blank and a PCR blank were included in each sequencing run (2 runs total). Due to limited DNA concentrations, we were unable to sequence 5 of the 6 PBS/Vehicle lung samples.

### Sequence processing and analysis

Primer removal and de-multiplexing was performed using Illumina Basespace using default settings. Sequence analysis was performed in R (72) with open-source software package ‘dada2’ (Version 1.16.0) (73). Each sequencing batch was processed separately until chimera removal. For each batch, the quality of the raw pair-end reads was visualized and used to determine appropriate truncation of read 1 (R1) by 10 bp and read 2 (R2) by 50 bp. After truncation, reads were discarded if they contained more than 2 maxEE “expected errors” or a quality score of less than or equal to 2. Following, each quality-filtered and trimmed read was processed independently by applying the trained dada2 algorithm. The reads were then merged with a minimum overlap of 20 bp. After merging, both sequencing batches that were previously processed separately were combined, and chimeras were removed using the consensus method with default settings. Taxonomy was assigned in ‘dada2’ using the native implementation of the naïve Bayesian classifier using Silva v.38 database. A count table and taxonomy file were created and used for downstream analysis.

Prior to diversity analysis, potential sequence contaminants were identified using package ‘decontam’ (Version 1.8.0) (74) in RStudio (Version 1.2.5042) (75). Briefly, potential contaminants were identified by using the prevalence-based contaminant identification, which relies on the principle that sequences from contaminating taxa have a higher prevalence in negative control samples (extraction and PCR blanks) than true samples (74). A threshold of 0.1 was used to identify contaminants. In total, 14 potential contaminants were identified by package ‘decontam’ (Table S1); however, all contaminants had biological relevance to the sample types collected except for one, Gemmobacter, which was removed from the data set.

Following, reads that did not identify as Bacteria, contained uncharacterized Phylum, identified as chloroplast and/or mitochondria were removed using the ‘phyloseq’ package (Version 1.32) (76). Subsequently, two samples with less than 10,000 reads/sample were removed (Lungs: Low/Vehicle at 2 dpc and High/Vehicle at 5 dpc) and one sample (Lungs: Mock/GC-376 at 14 dpc), considered an outlier according to Grubbs test on taxonomic abundance using the ‘outlier’ package (p = 6.022e-07) (77), was removed.

Alpha diversity metrics including observed number of amplicon sequence variants (ASVs), Shannon diversity and inverse Simpson (Inv Simpson) indexes were calculated using ‘phyloseq’. Briefly, samples were rarified to 12,000 using command *rrarefy* using the ‘vegan’ package (Version 2.57) (78). Following, the rarified counts were imported into ‘phyloseq’ and diversity indexes were calculated using command *estimaterichness*. Results were graphed using ‘ggplot2’ (Version 3.3.2) (79) and ‘ggpubr’ package (Version 0.4) (80). Statistical pair-wise comparison employing the Wilcox rank test was performed across groups and dpc. A Venn diagram of unique and shared ASVs was created using the package ‘microbiome’ (Version 1.10) (81). Rarified count data was converted to relative abundances, and then ASVs that were common among groups were combined. ASVs with a limited detection of 0.001 in at least 90% of the samples were included. The Venn diagram was graphed using package ‘eulerr’ (Version 6.1.0) (82). Regarding beta diversity, weighted Bray-Curtis dissimilarity matrix was calculated with a minimum of 20 and maximum of 100 random starts using the rarified count data in ‘vegan’. A Non-metric Multi-dimensional Scaling (NMDS) plot was used to graph the dissimilarity matrix using ‘ggplot2’. Ellipses were constructed using command *stat_ellipse* in ‘ggplot2’ with a multivariate t-distribution. All distances displayed in boxplots for comparison of within and across group Bray-Curtis dissimilarities were extracted from the same distance matrix as the one used for the NMDS and graphed using ‘ggplot2’. Hierarchal cluster analysis of the Bray-Curtis distances was created using command *hclust* with agglomeration method “average” (UPGMA) producing a cophenetic correlation coefficient of 0.79. The dendrogram was created using the function *plot* and shading/group colors were added using Adobe Illustrator (Version 25.0.1). Multivariate statistics was performed using permutational multivariate analysis of variance (PERMANOVA) tests using Bray-Curtis dissimilarity distances with 1000 permutations was generated using ‘vegan’ command *adonis2*. A p-value below 0.05 was considered significant. All other statistical tests were performed using Kruskal-Wallis or Wilcoxon signed-rank test using package ‘ggpubr’.

Relative abundances at the phylum and family level were generated using ‘phyloseq’. First, taxa were agglomerated at the phylum or family level and then transformed into relative abundance. Taxa that had less than 1% (ceca) or 2% (lung) abundance across all samples (separated by ceca and lungs) were grouped together. The box plots and bar plots of the relative abundances were generated using ‘ggplot2’. The three samples that were previously removed were not included in the analysis. Statistical pair-wise comparison among groups was performed using Wilcoxon signed-rank test. A p-value below 0.05 was considered significant. A rough estimation of Firmicutes/Bacteroidota ratio was calculated by dividing the relative abundance of the reads assigned to Firmicutes by the relative abundance of the reads assigned to Bacteroidota. Scripts used for analysis can be found on githubt at: https://github.com/brittanyaseibert/Seibertetal_SARS_K18hACE2Mice.

## Data Availability

The 16S sequencing dataset was deposited under BioProject PRJNA722991.

## ACKNOWLEDGMENTS

We thank the personnel from the Animal Health Research Center, University of Georgia. This study was supported by a subcontract from the Center for Research on Influenza Pathogenesis (CRIP) to D.R.P. under contract HHSN272201400008C from the National Institute of Allergy and Infectious Diseases (NIAID) Centers for Influenza Research and Surveillance (CEIRS). D.R.P. receives funds from the Georgia Research Alliance and the Caswell S. Eidson endowment fund, University of Georgia.

## SUPPLEMENTAL FIGURES

**Figure 1S. Alpha diversity metrics of ceca across time. (A)** Comparison of Observed ASVs, **(B)** Shannon diversity index and **(C)** Inverse Simpson of different groups (PBS/Vehicle Black, Low/Vehicle: Orange, High/Vehicle: Blue) separated by dpc from rarified ASV count table. Since the sample size was limited (n=2 or 3), pair-wise comparisons were not performed.

**Figure 2S. Alpha diversity metrics of cecum samples across time. (A)** Comparison of Observed ASVs, **(B)** Shannon diversity index and **(C)** Inverse Simpson of infected groups (Low/Vehicle and High/Vehicle) separated by dpc from rarified ASV count table. All statistical tests were performed using Kruskal-Wallis or Wilcox-rank test for pair-wise comparisons using * = p<0.05 ; ** = p<0.005; *** = p<0.0005; **** = p<0.00005.

**Figure 3S. Alpha diversity metrics of lung samples across time. (A)** Comparison of Observed ASVs, **(B)** Shannon diversity index and **(C)** Inverse Simpson of different groups (Mock/GC-376: Brown, Low/GC-376: Green, High/GC-376: Pink) separated by dpc from rarified ASV count table. Since the sample size was limited (n=2 or 3) pair-wise comparisons were not performed.

**Figure 4S. Alpha diversity metrics of lung samples across time. (A)** Comparison of Observed ASVs, **(B)** Shannon diversity index and **(C)** Inverse Simpson of infected groups (Low/GC-376 and High/GC-376) separated by dpc from rarified ASV count table. All statistical tests were performed using Kruskal-Wallis or Wilcox-rank test for pair-wise comparisons using * = p<0.05 ; ** = p<0.005; *** = p<0.0005; **** = p<0.00005.

**Figure 5S. Bray-Curtis Dissimilarity distances of lung samples without / with antiviral GC- 376**. Comparison of Bray-Curtis dissimilarity distances of mice infected with a A) low dose or (B) high dose of SARS-CoV-2 with or without GC-376. Gold boxes represent within-group variation while black boxes represent the between-group variation.

**TABLE S1.** Identification of Potential Contaminants by Decontam

| **ASV ID** | **Kingdom** | **Phylum** | **Class** | **Order** | **Family** | **Genus** |
| --- | --- | --- | --- | --- | --- | --- |
| ASV_249 | Bacteria | Firmicutes | Clostridia | Oscillospirales | Oscillospiraceae | Colidextribacter |
| ASV_314 | Bacteria | Bacteroidota | Bacteroidia | Flavobacteriales | Weeksellaceae | Empedobacter |
| ASV_355 | Bacteria | Firmicutes | Bacilli | Lactobacillales | Vagococcaceae | Vagococcus |
| ASV_395 | Bacteria | Proteobacteria | Gammaproteobacteria | Pseudomonadales | Moraxellaceae | Acinetobacter |
| ASV_398 | Bacteria | Proteobacteria | Gammaproteobacteria | Pseudomonadales | Moraxellaceae | Acinetobacter |
| ASV_470 | Bacteria | Proteobacteria | Alphaproteobacteria | Rhodobacterales | Rhodobacteraceae | Gemmobacter |
| ASV_522 | Bacteria | Firmicutes | Clostridia | Oscillospirales | Oscillospiraceae | NA |
| ASV_635 | Bacteria | Proteobacteria | Gammaproteobacteria | Pseudomonadales | Moraxellaceae | Acinetobacter |
| ASV_680 | Bacteria | Proteobacteria | Alphaproteobacteria | Sphingomonadales | Sphingomonadaceae | Sphingomonas |
| ASV_714 | Bacteria | Firmicutes | Bacilli | Lactobacillales | Streptococcaceae | Lactococcus |
| ASV_725 | Bacteria | Proteobacteria | Gammaproteobacteria | Pseudomonadales | Pseudomonadaceae | Pseudomonas |
| ASV_739 | Bacteria | Proteobacteria | Gammaproteobacteria | Burkholderiales | Comamonadaceae | NA |
| ASV_761 | Bacteria | Firmicutes | Clostridia | Oscillospirales | Oscillospiraceae | Flavonifractor |
| ASV_865 | Bacteria | Firmicutes | Clostridia | Lachnospirales | Lachnospiraceae | NA |

## REFERENCES

1. WHO. 2020. Coronavirus disease (COVID-19) pandemic. https://www.who.int/emergencies/diseases/novel-coronavirus-2019. Accessed

2. Hu B, Guo H, Zhou P, Shi ZL. 2020. Characteristics of SARS-CoV-2 and COVID-19. Nat Rev Microbiol doi:10.1038/s41579-020-00459-7.

3. Huang D, Lian X, Song F, Ma H, Lian Z, Liang Y, Qin T, Chen W, Wang S. 2020. Clinical features of severe patients infected with 2019 novel coronavirus: a systematic review and meta-analysis. Ann Transl Med 8:576.

4. Mehta P, McAuley DF, Brown M, Sanchez E, Tattersall RS, Manson JJ, Hlh Across Speciality Collaboration UK. 2020. COVID-19: consider cytokine storm syndromes and immunosuppression. Lancet 395:1033–1034.

5. Gu S, Chen Y, Wu Z, Chen Y, Gao H, Lv L, Guo F, Zhang X, Luo R, Huang C, Lu H, Zheng B, Zhang J, Yan R, Zhang H, Jiang H, Xu Q, Guo J, Gong Y, Tang L, Li L. 2020. Alterations of the Gut Microbiota in Patients with COVID-19 or H1N1 Influenza. Clin Infect Dis doi:10.1093/cid/ciaa709.

6. Jin X, Lian JS, Hu JH, Gao J, Zheng L, Zhang YM, Hao SR, Jia HY, Cai H, Zhang XL, Yu GD, Xu KJ, Wang XY, Gu JQ, Zhang SY, Ye CY, Jin CL, Lu YF, Yu X, Yu XP, Huang JR, Xu KL, Ni Q, Yu CB, Zhu B, Li YT, Liu J, Zhao H, Zhang X, Yu L, Guo YZ, Su JW, Tao JJ, Lang GJ, Wu XX, Wu WR, Qv TT, Xiang DR, Yi P, Shi D, Chen Y, Ren Y, Qiu YQ, Li LJ, Sheng J, Yang Y. 2020. Epidemiological, clinical and virological characteristics of 74 cases of coronavirus-infected disease 2019 (COVID-19) with gastrointestinal symptoms. Gut 69:1002–1009.

7. Du M, Cai G, Chen F, Christiani DC, Zhang Z, Wang M. 2020. Multiomics Evaluation of Gastrointestinal and Other Clinical Characteristics of COVID-19. Gastroenterology 158:2298–2301 e7.

8. Qian Q, Fan L, Liu W, Li J, Yue J, Wang M, Ke X, Yin Y, Chen Q, Jiang C. 2020. Direct evidence of active SARS-CoV-2 replication in the intestine. Clin Infect Dis doi:10.1093/cid/ciaa925.

9. Chen Y, Chen L, Deng Q, Zhang G, Wu K, Ni L, Yang Y, Liu B, Wang W, Wei C, Yang J, Ye G, Cheng Z. 2020. The presence of SARS-CoV-2 RNA in the feces of COVID-19 patients. J Med Virol 92:833–840.

10. Conte L, Toraldo DM. 2020. Targeting the gut-lung microbiota axis by means of a high-fibre diet and probiotics may have anti-inflammatory effects in COVID-19 infection. Ther Adv Respir Dis 14:1753466620937170.

11. Chen N, Zhou M, Dong X, Qu J, Gong F, Han Y, Qiu Y, Wang J, Liu Y, Wei Y, Xia J, Yu T, Zhang X, Zhang L. 2020. Epidemiological and clinical characteristics of 99 cases of 2019 novel coronavirus pneumonia in Wuhan, China: a descriptive study. Lancet 395:507–513.

12. Wang D, Hu B, Hu C, Zhu F, Liu X, Zhang J, Wang B, Xiang H, Cheng Z, Xiong Y, Zhao Y, Li Y, Wang X, Peng Z. 2020. Clinical Characteristics of 138 Hospitalized Patients With 2019 Novel Coronavirus-Infected Pneumonia in Wuhan, China. JAMA 323:1061–1069.

13. Pichon M, Lina B, Josset L. 2017. Impact of the Respiratory Microbiome on Host Responses to Respiratory Viral Infection. Vaccines (Basel) 5.

14. Chen CJ, Wu GH, Kuo RL, Shih SR. 2017. Role of the intestinal microbiota in the immunomodulation of influenza virus infection. Microbes Infect 19:570–579.

15. Bharti R, Grimm DG. 2021. Current challenges and best-practice protocols for microbiome analysis. Brief Bioinform 22:178–193.

16. Gu L, Deng H, Ren Z, Zhao Y, Yu S, Guo Y, Dai J, Chen X, Li K, Li R, Wang G. 2019. Dynamic Changes in the Microbiome and Mucosal Immune Microenvironment of the Lower Respiratory Tract by Influenza Virus Infection. Front Microbiol 10:2491.

17. Rowe HM, Livingston B, Margolis E, Davis A, Meliopoulos VA, Echlin H, Schultz-Cherry S, Rosch JW. 2020. Respiratory Bacteria Stabilize and Promote Airborne Transmission of Influenza A Virus. mSystems 5.

18. Yildiz S, Mazel-Sanchez B, Kandasamy M, Manicassamy B, Schmolke M. 2018. Influenza A virus infection impacts systemic microbiota dynamics and causes quantitative enteric dysbiosis. Microbiome 6:9.

19. Deriu E, Boxx GM, He X, Pan C, Benavidez SD, Cen L, Rozengurt N, Shi W, Cheng G. 2016. Influenza Virus Affects Intestinal Microbiota and Secondary Salmonella Infection in the Gut through Type I Interferons. PLoS Pathog 12:e1005572.

20. Groves HT, Cuthbertson L, James P, Moffatt MF, Cox MJ, Tregoning JS. 2018. Respiratory Disease following Viral Lung Infection Alters the Murine Gut Microbiota. Front Immunol 9:182.

21. Sencio V, Barthelemy A, Tavares LP, Machado MG, Soulard D, Cuinat C, Queiroz-Junior CM, Noordine ML, Salome-Desnoulez S, Deryuter L, Foligne B, Wahl C, Frisch B, Vieira AT, Paget C, Milligan G, Ulven T, Wolowczuk I, Faveeuw C, Le Goffic R, Thomas M, Ferreira S, Teixeira MM, Trottein F. 2020. Gut Dysbiosis during Influenza Contributes to Pulmonary Pneumococcal Superinfection through Altered Short-Chain Fatty Acid Production. Cell Rep 30:2934–2947 e6.

22. Hu X, Zhao Y, Yang Y, Gong W, Sun X, Yang L, Zhang Q, Jin M. 2020. Akkermansia muciniphila Improves Host Defense Against Influenza Virus Infection. Front Microbiol 11:586476.

23. Oh JZ, Ravindran R, Chassaing B, Carvalho FA, Maddur MS, Bower M, Hakimpour P, Gill KP, Nakaya HI, Yarovinsky F, Sartor RB, Gewirtz AT, Pulendran B. 2014. TLR5-mediated sensing of gut microbiota is necessary for antibody responses to seasonal influenza vaccination. Immunity 41:478–492.

24. Zuo T, Zhang F, Lui GCY, Yeoh YK, Li AYL, Zhan H, Wan Y, Chung ACK, Cheung CP, Chen N, Lai CKC, Chen Z, Tso EYK, Fung KSC, Chan V, Ling L, Joynt G, Hui DSC, Chan FKL, Chan PKS, Ng SC. 2020. Alterations in Gut Microbiota of Patients With COVID-19 During Time of Hospitalization. Gastroenterology 159:944–955 e8.

25. Yang T, Chakraborty S, Piu Saha BM, Cheng X, Yeo J-Y, Mei X, Zhou G, Mandal J, Golonka R, Yeoh BS, Putluri V, Piyarathna DWB, Putluri N, McCarthy CG, Wenceslau CF, Sreekumar A, Gewirtz AT, Vijay-Kumar M, Joe B. 2020. Gnotobiotic Rats Reveal That Gut Microbiota Regulates Colonic mRNA of Ace2, the Receptor for SARS-CoV-2 Infectivity. Hypertension 76:1–3.

26. Zuo T, Liu Q, Zhang F, Lui GC, Tso EY, Yeoh YK, Chen Z, Boon SS, Chan FK, Chan PK, Ng SC. 2020. Depicting SARS-CoV-2 faecal viral activity in association with gut microbiota composition in patients with COVID-19. Gut doi:10.1136/gutjnl-2020-322294.

27. Mostafa HH, Fissel JA, Fanelli B, Bergman Y, Gniazdowski V, Dadlani M, Carroll KC, Colwell RR, Simner PJ. 2020. Metagenomic Next-Generation Sequencing of Nasopharyngeal Specimens Collected from Confirmed and Suspect COVID-19 Patients. mBio 11.

28. De Maio F, Posteraro B, Ponziani FR, Cattani P, Gasbarrini A, Sanguinetti M. 2020. Nasopharyngeal Microbiota Profiling of SARS-CoV-2 Infected Patients. Biol Proced Online 22:18.

29. Cheung KS, Hung IFN, Chan PPY, Lung KC, Tso E, Liu R, Ng YY, Chu MY, Chung TWH, Tam AR, Yip CCY, Leung KH, Fung AY, Zhang RR, Lin Y, Cheng HM, Zhang AJX, To KKW, Chan KH, Yuen KY, Leung WK. 2020. Gastrointestinal Manifestations of SARS-CoV-2 Infection and Virus Load in Fecal Samples From a Hong Kong Cohort: Systematic Review and Meta-analysis. Gastroenterology 159:81–95.

30. Ceccarelli G, Borrazzo C, Pinacchio C, Santinelli L, Innocenti GP, Cavallari EN, Celani L, Marazzato M, Alessandri F, Ruberto F, Pugliese F, Venditti M, Mastroianni CM, d’Ettorre G. 2021. Oral Bacteriotherapy in Patients With COVID-19: A Retrospective Cohort Study. Front Nutr 7:613928.

31. Durban A, Abellan JJ, Jimenez-Hernandez N, Ponce M, Ponce J, Sala T, D’Auria G, Latorre A, Moya A. 2011. Assessing gut microbial diversity from feces and rectal mucosa. Microb Ecol 61:123–33.

32. Ingala MR, Simmons NB, Wultsch C, Krampis K, Speer KA, Perkins SL. 2018. Comparing Microbiome Sampling Methods in a Wild Mammal: Fecal and Intestinal Samples Record Different Signals of Host Ecology, Evolution. Front Microbiol 9:803.

33. Kozik AJ, Nakatsu CH, Chun H, Jones-Hall YL. 2019. Comparison of the fecal, cecal, and mucus microbiome in male and female mice after TNBS-induced colitis. PLoS One 14:e0225079.

34. Kolde R, Franzosa EA, Rahnavard G, Hall AB, Vlamakis H, Stevens C, Daly MJ, Xavier RJ, Huttenhower C. 2018. Host genetic variation and its microbiome interactions within the Human Microbiome Project. Genome Med 10:6.

35. Voigt AY, Costea PI, Kultima JR, Li SS, Zeller G, Sunagawa S, Bork P. 2015. Temporal and technical variability of human gut metagenomes. Genome Biol 16:73.

36. Karl JP, Hatch AM, Arcidiacono SM, Pearce SC, Pantoja-Feliciano IG, Doherty LA, Soares JW. 2018. Effects of Psychological, Environmental and Physical Stressors on the Gut Microbiota. Front Microbiol 9:2013.

37. Spor A, Koren O, Ley R. 2011. Unravelling the effects of the environment and host genotype on the gut microbiome. Nat Rev Microbiol 9:279–90.

38. Moreau GB, Burgess SL, Sturek JM, Donlan AN, Petri WA, Mann BJ. 2020. Evaluation of K18-hACE2 Mice as a Model of SARS-CoV-2 Infection. Am J Trop Med Hyg 103:1215–1219.

39. Zheng J, Roy Wong LY, Li K, Verma AK, Ortiz M, Wohlford-Lenane C, Leidinger MR, Knudson CM, Meyerholz DK, McCray PB, Perlman S. 2020. K18-hACE2 Mice for Studies of COVID-19 Treatments and Pathogenesis Including Anosmia. bioRxiv doi:10.1101/2020.08.07.242073.

40. Rathnasinghe R, Strohmeier S, Amanat F, Gillespie VL, Krammer F, Garcia-Sastre A, Coughlan L, Schotsaert M, Uccellini MB. 2020. Comparison of transgenic and adenovirus hACE2 mouse models for SARS-CoV-2 infection. Emerg Microbes Infect 9:2433–2445.

41. Winkler ES, Bailey AL, Kafai NM, Nair S, McCune BT, Yu J, Fox JM, Chen RE, Earnest JT, Keeler SP, Ritter JH, Kang LI, Dort S, Robichaud A, Head R, Holtzman MJ, Diamond MS. 2020. Publisher Correction: SARS-CoV-2 infection of human ACE2-transgenic mice causes severe lung inflammation and impaired function. Nat Immunol 21:1470.

42. Oladunni FS, Park JG, Pino PA, Gonzalez O, Akhter A, Allue-Guardia A, Olmo-Fontanez A, Gautam S, Garcia-Vilanova A, Ye C, Chiem K, Headley C, Dwivedi V, Parodi LM, Alfson KJ, Staples HM, Schami A, Garcia JI, Whigham A, Platt RN, 2nd, Gazi M, Martinez J, Chuba C, Earley S, Rodriguez OH, Mdaki SD, Kavelish KN, Escalona R, Hallam CRA, Christie C, Patterson JL, Anderson TJC, Carrion R, Jr., Dick EJ, Jr., Hall-Ursone S, Schlesinger LS, Alvarez X, Kaushal D, Giavedoni LD, Turner J, Martinez-Sobrido L, Torrelles JB. 2020. Lethality of SARS-CoV-2 infection in K18 human angiotensin-converting enzyme 2 transgenic mice. Nat Commun 11:6122.

43. Cáceres CJ, Cardenas-Garcia S, Carnaccini S, Seibert B, Rajao DS, Wang J, Perez DR. Accepted. Efficacy of GC-376 against SARS-CoV-2 virus infection in the K18 hACE2 transgenic mouse model. Scientific Reports.

44. Golden JW, Cline CR, Zeng X, Garrison AR, Carey BD, Mucker EM, White LE, Shamblin JD, Brocato RL, Liu J, Babka AM, Rauch HB, Smith JM, Hollidge BS, Fitzpatrick C, Badger CV, Hooper JW. 2020. Human angiotensin-converting enzyme 2 transgenic mice infected with SARS-CoV-2 develop severe and fatal respiratory disease. JCI Insight 5.

45. Zheng J, Wong LR, Li K, Verma AK, Ortiz ME, Wohlford-Lenane C, Leidinger MR, Knudson CM, Meyerholz DK, McCray PB, Jr., Perlman S. 2020. COVID-19 treatments and pathogenesis including anosmia in K18-hACE2 mice. Nature doi:10.1038/s41586-020-2943-z.

46. Yinda CK, Port JR, Bushmaker T, Offei Owusu I, Purushotham JN, Avanzato VA, Fischer RJ, Schulz JE, Holbrook MG, Hebner MJ, Rosenke R, Thomas T, Marzi A, Best SM, de Wit E, Shaia C, van Doremalen N, Munster VJ. 2021. K18-hACE2 mice develop respiratory disease resembling severe COVID-19. PLoS Pathog 17:e1009195.

47. Cao J, Wang C, Zhang Y, Lei G, Xu K, Zhao N, Lu J, Meng F, Yu L, Yan J, Bai C, Zhang S, Zhang N, Gong Y, Bi Y, Shi Y, Chen Z, Dai L, Wang J, Yang P. 2021. Integrated gut virome and bacteriome dynamics in COVID-19 patients. Gut Microbes 13:1–21.

48. Yinda CK, Port JR, Bushmaker T, Owusu IO, Avanzato VA, Fischer RJ, Schulz JE, Holbrook MG, Hebner MJ, Rosenke R, Thomas T, Marzi A, Best SM, de Wit E, Shaia C, van Doremalen N, Munster VJ. 2020. K18-hACE2 mice develop respiratory disease resembling severe COVID-19. bioRxiv doi:10.1101/2020.08.11.246314.

49. Kalantar-Zadeh K, Ward SA, Kalantar-Zadeh K, El-Omar EM. 2020. Considering the Effects of Microbiome and Diet on SARS-CoV-2 Infection: Nanotechnology Roles. ACS Nano 14:5179–5182.

50. Wilks J, Beilinson H, Golovkina TV. 2013. Dual role of commensal bacteria in viral infections. Immunol Rev 255:222–9.

51. Kuss SK, Best GT, Etheredge CA, Pruijssers AJ, Frierson JM, Hooper LV, Dermody TS, Pfeiffer JK. 2011. Intestinal microbiota promote enteric virus replication and systemic pathogenesis. Science 334:249–52.

52. Robinson CM, Jesudhasan PR, Pfeiffer JK. 2014. Bacterial lipopolysaccharide binding enhances virion stability and promotes environmental fitness of an enteric virus. Cell Host Microbe 15:36–46.

53. Robinson CM. 2019. Enteric viruses exploit the microbiota to promote infection. Curr Opin Virol 37:58–62.

54. Robinson CM, Pfeiffer JK. 2014. Viruses and the Microbiota. Annu Rev Virol 1:55–69.

55. Stojanov S, Berlec A, Strukelj B. 2020. The Influence of Probiotics on the Firmicutes/Bacteroidetes Ratio in the Treatment of Obesity and Inflammatory Bowel disease. Microorganisms 8.

56. Vacca M, Celano G, Calabrese FM, Portincasa P, Gobbetti M, De Angelis M. 2020. The Controversial Role of Human Gut Lachnospiraceae. Microorganisms 8.

57. Bartley JM, Zhou X, Kuchel GA, Weinstock GM, Haynes L. 2017. Impact of Age, Caloric Restriction, and Influenza Infection on Mouse Gut Microbiome: An Exploratory Study of the Role of Age-Related Microbiome Changes on Influenza Responses. Front Immunol 8:1164.

58. Hadjadj J, Yatim N, Barnabei L, Corneau A, Boussier J, Smith N, Pere H, Charbit B, Bondet V, Chenevier-Gobeaux C, Breillat P, Carlier N, Gauzit R, Morbieu C, Pene F, Marin N, Roche N, Szwebel TA, Merkling SH, Treluyer JM, Veyer D, Mouthon L, Blanc C, Tharaux PL, Rozenberg F, Fischer A, Duffy D, Rieux-Laucat F, Kerneis S, Terrier B. 2020. Impaired type I interferon activity and inflammatory responses in severe COVID-19 patients. Science 369:718–724.

59. Hanada S, Pirzadeh M, Carver KY, Deng JC. 2018. Respiratory Viral Infection-Induced Microbiome Alterations and Secondary Bacterial Pneumonia. Front Immunol 9:2640.

60. Geerlings SY, Kostopoulos I, de Vos WM, Belzer C. 2018. Akkermansia muciniphila in the Human Gastrointestinal Tract: When, Where, and How? Microorganisms 6.

61. Derrien M, Van Baarlen P, Hooiveld G, Norin E, Muller M, de Vos WM. 2011. Modulation of Mucosal Immune Response, Tolerance, and Proliferation in Mice Colonized by the Mucin-Degrader Akkermansia muciniphila. Front Microbiol 2:166.

62. LeMessurier KS, Iverson AR, Chang TC, Palipane M, Vogel P, Rosch JW, Samarasinghe AE. 2019. Allergic inflammation alters the lung microbiome and hinders synergistic co-infection with H1N1 influenza virus and Streptococcus pneumoniae in C57BL/6 mice. Sci Rep 9:19360.

63. Le Noci V, Guglielmetti S, Arioli S, Camisaschi C, Bianchi F, Sommariva M, Storti C, Triulzi T, Castelli C, Balsari A, Tagliabue E, Sfondrini L. 2018. Modulation of Pulmonary Microbiota by Antibiotic or Probiotic Aerosol Therapy: A Strategy to Promote Immunosurveillance against Lung Metastases. Cell Rep 24:3528–3538.

64. Ma ZS. 2020. Testing the Anna Karenina Principle in Human Microbiome-Associated Diseases. iScience 23:101007.

65. Zaneveld JR, McMinds R, Vega Thurber R. 2017. Stress and stability: applying the Anna Karenina principle to animal microbiomes. Nat Microbiol 2:17121.

66. Khatiwada S, Subedi A. 2020. Lung microbiome and coronavirus disease 2019 (COVID-19): Possible link and implications. Hum Microb J 17:100073.

67. Gollwitzer ES, Saglani S, Trompette A, Yadava K, Sherburn R, McCoy KD, Nicod LP, Lloyd CM, Marsland BJ. 2014. Lung microbiota promotes tolerance to allergens in neonates via PD-L1. Nat Med 20:642–7.

68. Leung RK, Zhou JW, Guan W, Li SK, Yang ZF, Tsui SK. 2013. Modulation of potential respiratory pathogens by pH1N1 viral infection. Clin Microbiol Infect 19:930–5.

69. Sharma NS, Vestal G, Wille K, Patel KN, Cheng F, Tipparaju S, Tousif S, Banday MM, Xu X, Wilson L, Nair VS, Morrow C, Hayes D, Jr., Seyfang A, Barnes S, Deshane JS, Gaggar A. 2020. Differences in airway microbiome and metabolome of single lung transplant recipients. Respir Res 21:104.

70. Reed LJ, Muench H. 1938. A simple method for estimating fifty percent endpoints. Am J Hyg 27:493–497.

71. Caporaso JG, Lauber CL, Walters WA, Berg-Lyons D, Huntley J, Fierer N, Owens SM, Betley J, Fraser L, Bauer M, Gormley N, Gilbert JA, Smith G, Knight R. 2012. Ultra-high-throughput microbial community analysis on the Illumina HiSeq and MiSeq platforms. ISME J 6:1621–4.

72. Team RC. 2014. R: A language and environment for statistical computing, R Foundation for Statistical Computing, http://www.R-project.org/.

73. Callahan BJ, McMurdie PJ, Rosen MJ, Han AW, Johnson AJ, Holmes SP. 2016. DADA2: High-resolution sample inference from Illumina amplicon data. Nat Methods 13:581–3.

74. Davis NM, Proctor DM, Holmes SP, Relman DA, Callahan BJ. 2018. Simple statistical identification and removal of contaminant sequences in marker-gene and metagenomics data. Microbiome 6:226.

75. Team R. 2020. RStudio: Integrated Development for R, PBC, Boston, MA. http://www.rstudio.com/.

76. McMurdie PJ, Holmes S. 2013. phyloseq: an R package for reproducible interactive analysis and graphics of microbiome census data. PLoS One 8:e61217.

77. Grubbs FE. 1950. Sample Criteria for testing outlying observations. Ann Math Stat 21:27–58.

78. Jari Oksanen, F. Guillaume Blanchet, Michael Friendly, Roeland Kindt, Pierre Legendre, Dan McGlinn, Peter R. Minchin, R. B. O’Hara, Gavin L. Simpson, Peter Solymos, M. Henry H. Stevens, Eduard Szoecs, Wagner H. 2020. vegan: Community Ecology Package, vR package version 2.5-7. https://CRAN.R-project.org/package=vegan.

79. Wickham H. 2016. ggplot2: Elegant Graphics for Data, Springer-Verlag, New York. https://ggplot2.tidyverse.org.

80. Kassambara A. 2020. ggpubr: ‘ggplot2’ Based Publication Ready Plots, vR package version 0.4.0. https://CRAN.R-project.org/package=ggpubr.

81. Leo Lahti, Shetty S. 2012-2019. microbiome R package, http://microbiome.github.io.

82. Larsson J. 2020. _eulerr: Area-Proportional Euler and Venn Diagrams with Ellipses_, vR package version 6.1.0. https://cran.r-project.org/package=eulerr.

